# DIET INFLUENCES PERIPHERAL AMYLOID β METABOLISM: A ROLE FOR CIRCULATING INSULIN-LIKE GROWTH FACTOR I

**DOI:** 10.1101/2020.07.09.180265

**Authors:** R. Herrero-Labrador, A. Trueba-Saiz, L. Martinez-Rachadell, E. Fernandez de Sevilla, S. Diaz-Pacheco, Ana F. Fernandez, I. Torres Aleman

## Abstract

Obesity is a risk factor for Alzheimer’s disease (AD), but underlying mechanisms are not clear. We analyzed peripheral clearance of amyloid β (Aβ) in overweight mice because its systemic elimination may impact on brain Aβ load, a major landmark of AD pathology. Overweight mice showed increased peripheral Aβ clearance by the liver, the major site of elimination of systemic Aβ, but unaltered brain Aβ levels. Since circulating insulin-like growth factor I (IGF-I) modulates brain Aβ clearance, and is increased in serum of overweight mice, we determined whether it affects peripheral Aβ clearance. We found that Aβ uptake by hepatocytes is stimulated by IGF-I. Moreover, mice with low serum IGF-I levels show reduced peripheral Aβ clearance. In the brain, IGF-I favored association of its receptor (IGF-IR) with Aβ precursor protein (APP), and at the same time stimulated non-amyloidogenic processing of APP in astrocytes, as indicated by an increased sAPPα/sAPPβ ratio after IGF-I treatment. Since serum IGF-I enters into the brain in an activity-dependent manner, we analyzed in overweight mice the effect of brain activation by environmental enrichment (EE) on brain IGF-IR phosphorylation and its association to APP, as a readout of IGF-I activity. After EE, significantly less activation of brain IGF-IR phosphorylation and APP/IGF-IR association was found in overweight mice as compared to lean controls. Collectively, these results indicate that diet influences peripheral clearance of Aβ without affecting brain Aβ load. Increased serum IGF-I likely contributes to enhanced peripheral Aβ clearance in overweight mice, without affecting brain Aβ clearance probably because its brain entrance is reduced.

## Introduction

Obesity is considered a risk factor for AD (1-3). However, the relationship between body weight and dementia appears complex (4-6), and recent observations even pose a protective role of late-life excess weight in AD (7). Taking into account the worrying worldwide prevalence of obesity and dementia (8,9), greater knowledge of possible links between the two conditions is imperative. Amyloid β (Aβ) handling may be one such link, as this peptide is considered a major pathogenic factor in AD and obesity-associated inflammation (10) may interfere with it.

We recently proposed that insulin peptides, including insulin and insulin-like growth factor I (IGF-I), may be involved in the connection between life-style and AD risk (11), although apparently contradictory evidence links IGF-I with AD (12,13). Significantly, the actions of IGF-I on the brain are modulated by diet (14). Brain IGF-I is in part locally synthesized (15), and in part derived from uptake from the circulation (16). The entrance of circulating IGF-I into the brain is tightly regulated (17), probably because it participates in many essential brain functions (18).

IGF-I may not only be involved in brain Aβ clearance (12); other findings point to an effect of IGF-I on APP processing towards the non-amyloidogenic pathway, reducing in this way its production (19-21). However, IGF-I has also been shown to favor the amyloidogenic pathway (22,23), while a deleterious effect of IGF-I signaling in proteostasis, favoring Aβ accumulation, has also been reported (13). Of note, hepatocytes, the main source of circulating IGF-I (24), are the major disposal system for circulating Aβ in mice (25), and previous evidence has shown that insulin, a hormone closely related to IGF-I, favors Aβ uptake by hepatocytes (26).

In the present work we investigated regulation of peripheral Aβ clearance in overweight mice, its impact on brain Aβ levels, and the role of circulating IGF-I.

## Results

### Diet influences peripheral Aβ clearance

We examined peripheral Aβ disposal in overweight mice because is a proposed mechanism for central Aβ clearance (27). We administered fluorescently tagged Aβ to mice fed with a high fat diet (HFD) for 10 weeks. Animals became overweight and glucose intolerant (Suppl Figure A,B). Ninety minutes after intravenous injection of Aβ, overweight mice showed significantly increased fluorescence accumulation in the liver and decreased in serum, suggesting increased disposal of Aβ through hepatocytes (Figure 1A). Conversely, brain Aβ levels in overweight mice were not different from those seen in lean ones (Figure 1B).

**Figure 1.**
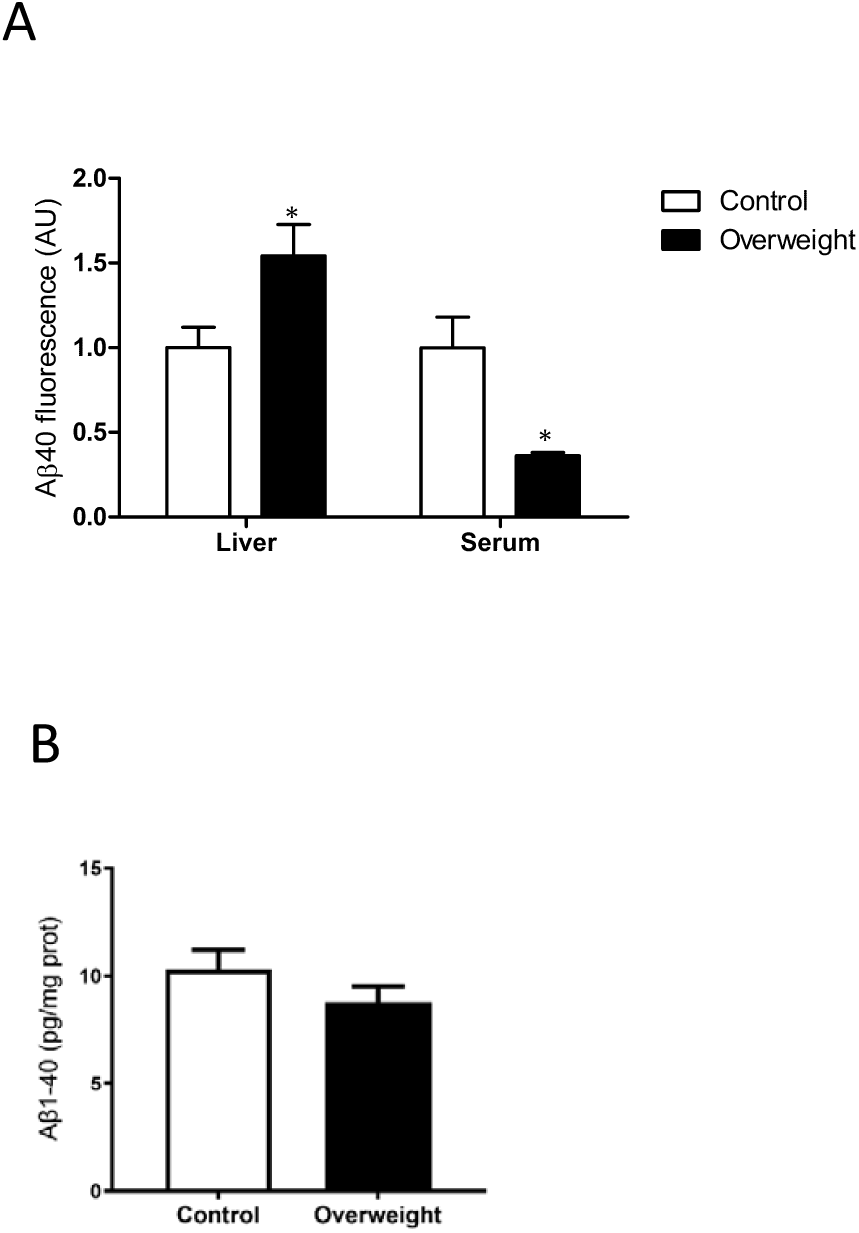
Diet modulates peripheral Aβ uptake. **A**, Overweight mice show enhanced Aβ uptake by the liver and lower serum Aβ levels (n=4). **B**, Brain Aβ levels remain unaltered in overweight mice (n=6-8). *p<0.05, **p<0.01, and ***p<0.001 in this and following figures.

### IGF-I promotes Aβ uptake by hepatocytes

To try to clarify the discrepancy between increased peripheral disposal of Aβ and normal brain Aβ load, we analyzed a possible role of IGF-I, that is mostly produced by the liver (24), and is increased in overweight mice (Figure 2A). We determined whether IGF-I modulates uptake of Aβ by hepatocytes. As shown in Figure 2B, in the presence of IGF-I (10 nM), hepatocytes accumulated significantly more fluorescence, suggesting a stimulatory action of IGF-I on Aβ uptake by these cells. Moreover, liver IGF-I deficient (LID) mice with a 70% reduction in circulating IGF-I (28), showed reduced liver accumulation of tagged Aβ after intravenous injection, while blood levels were increased, as compared to controls, indicating reduced liver clearance (Figure 2C). Collectively, this suggests that serum IGF-I modulates liver clearance of Aβ.

**Figure 2.**
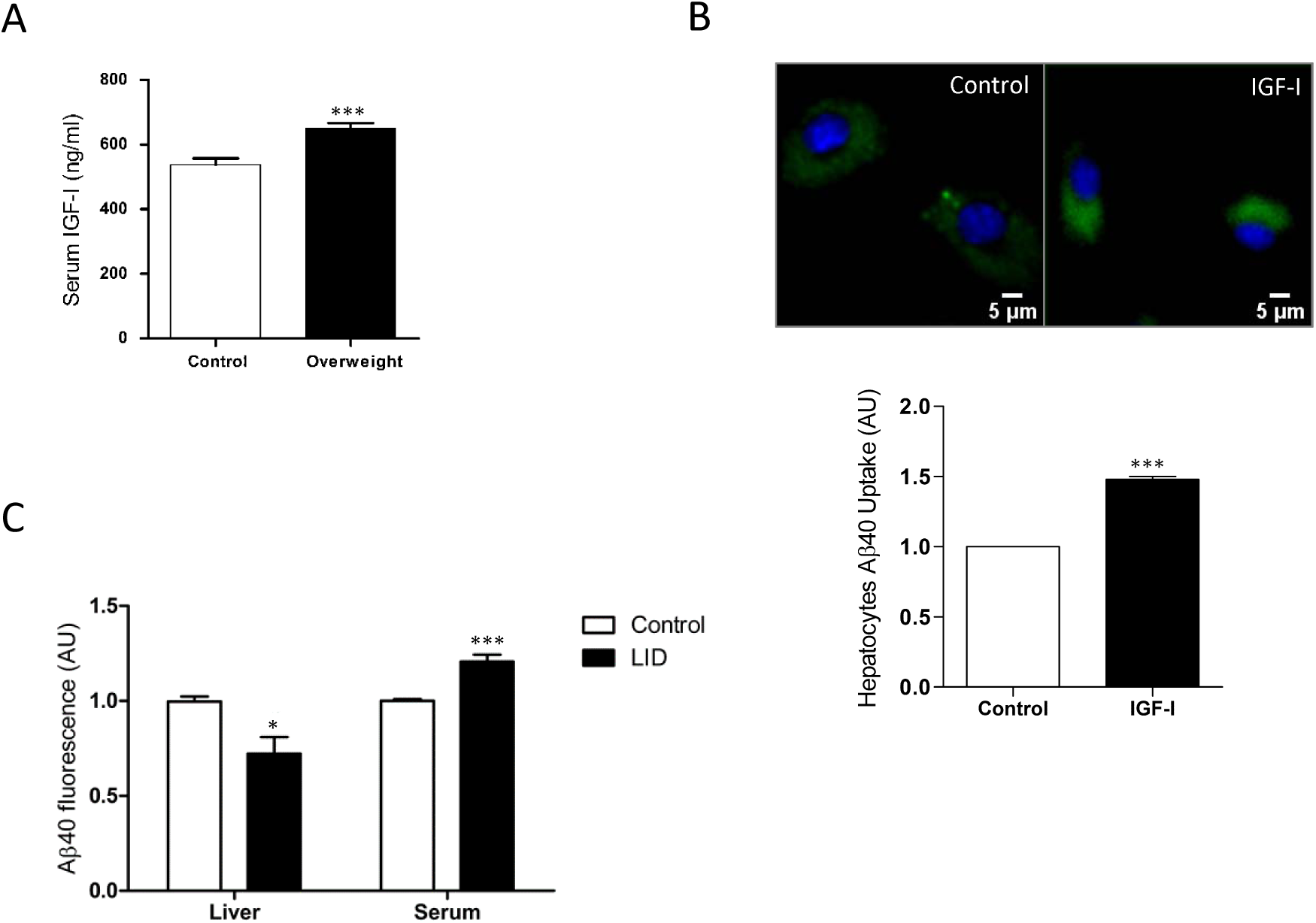
Modulation by IGF-I of Aβ uptake by hepatocytes. **A**, Serum levels of IGF-I are increased after 10 weeks of high fat diet (n=10 per group). **B**, IGF-I induces uptake of Aβ by hepatocytes (n= 6). Representative micrograph of cultured hepatocytes with internalized fluorescent Aβ (green). Cell nuclei stained with Hoescht. Lower histograms: quantification of intracellular fluorescent Aβ after IGF-I treatment. **C**, Serum IGF-I deficient mice (LID mice) show reduced Aβ uptake by the liver (n= 5 control/6 LID).

### Cell-specific actions of IGF-I in APP metabolism by brain cells

IGF-I has been reported to promote either amyloidogenic (22), or non-amyloidogenic (20) APP processing pathways in neuronal cell lines. To clarify its role in primary cells, we analyzed the actions of IGF-I on amyloidogenic and non-amyloidogenic APP processing by astrocytes and neurons, the primary sources of Aβ in the brain (29,30). Using the soluble APP metabolites sAPPβ and sAPPα as markers of the amyloidogenic and the non-amyloidogenic pathway, respectively, we found that IGF-I modulates their production in a cell specific fashion. In astrocytes, secretion of both soluble forms of APP was stimulated by IGF-I, whereas in neurons IGF-I inhibited their secretion (Figure 3A). However, the APPα/sAPPβ ratio was increased in both cell types, indicating that the net action of IGF-I is to promote non-amyloidogenic processing of APP (Figure 3B).

**Figure 3.**
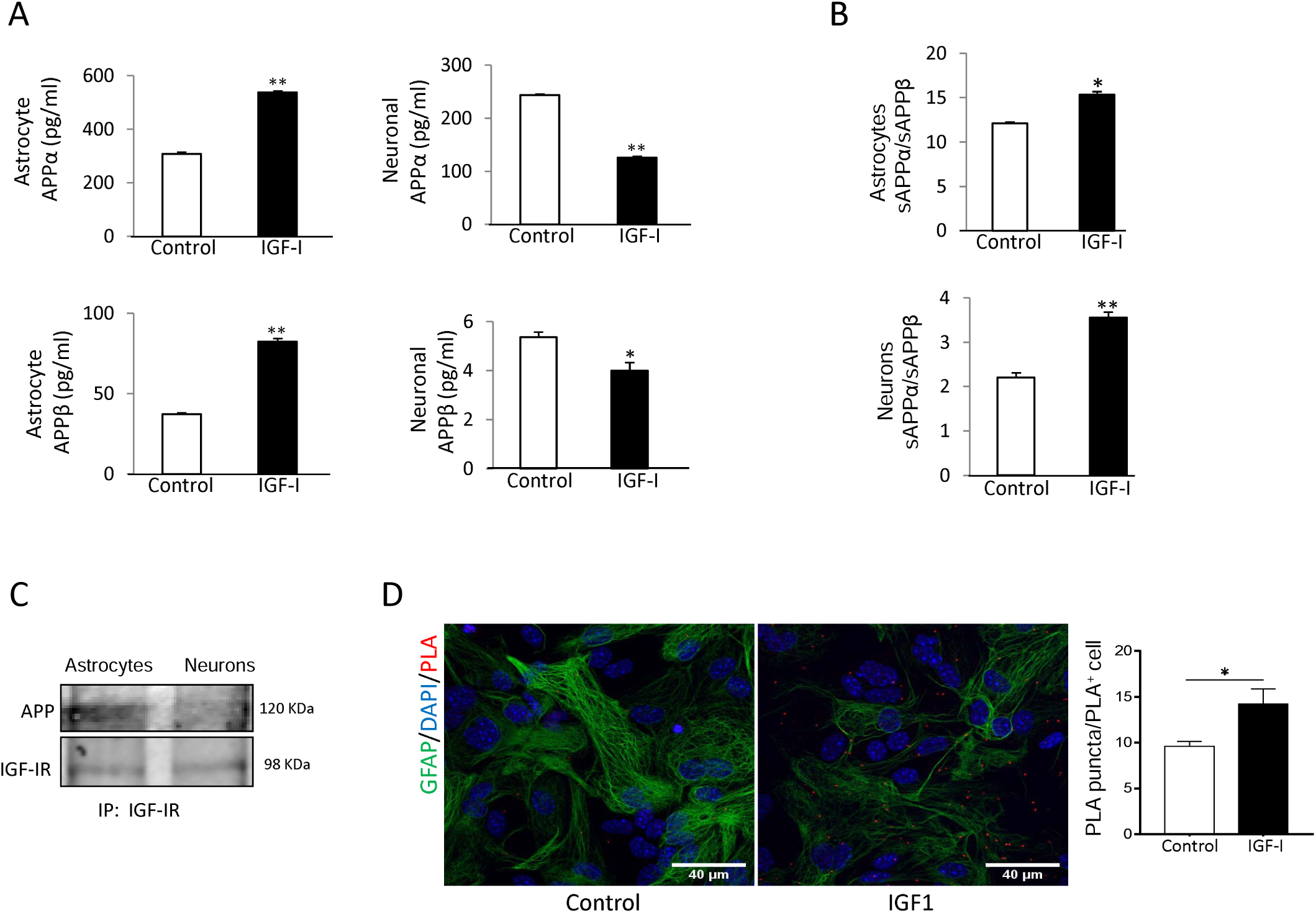
APP processing in astrocytes is modulated by IGF-I. **A**, IGF-I stimulated the secretion of both sAPPα and sAPPβ in cultured astrocytes (left histograms), while inhibited it in neurons (righ histograms, n=4). **B**, However, IGF-I increased the sAPPα/sAPPβ ratio in both cell types, indicating a net non-amyloidogenic action of IGF-I in these cells. **C**, IGF-IR and APP co-immunoprecipitate in cultured astrocytes while in neurons the interaction is negligible. **D**, Proximity ligation assays (PLA) of APP and IGF-IR in cultured astrocytes confirm an interaction of both proteins that is upregulated by IGF-I (n= 3). Cell nuclei stained with Hoescht.

Since both IGF-IR and APP associate to LRP1, and APP processing depends on its subcellular localization (31), we assessed whether IGF-IR and APP interact with each other. Indeed, IGF-IR and APP co-immunoprecipitated in astrocytes, whereas in neurons the interaction was negligible (Figure 3C). Proximity ligation assays (PLA) confirmed a robust interaction of APP with IGF-IR in astrocytes (Figure 3D), while in neurons the interaction was negligible (not shown). Treatment of astrocytes with IGF-I resulted in a significantly increased interaction between both proteins, as determined by a stronger PLA signal (Figure 3D).

Since IGF-I promotes Aβ uptake by hepatocytes (32), we examined whether it can exert similar action in brain cells. In this organ, the main cell types involved in Aβ clearance are microglia and astrocytes through its uptake and degradation (33,34), and endothelial cells at the blood-brain-barrier (BBB), through efflux of brain Aβ into the circulation (35). We found that IGF-I promoted Aβ uptake by astrocytes (Figure 4A), while decreased it in microglia (Figure 4B). In brain endothelial cell cultures mimicking the BBB architecture (17), IGF-I not significantly inhibited Aβ efflux from the “brain” side to the “blood” side of the double chamber (Figure 4C).

**Figure 4.**
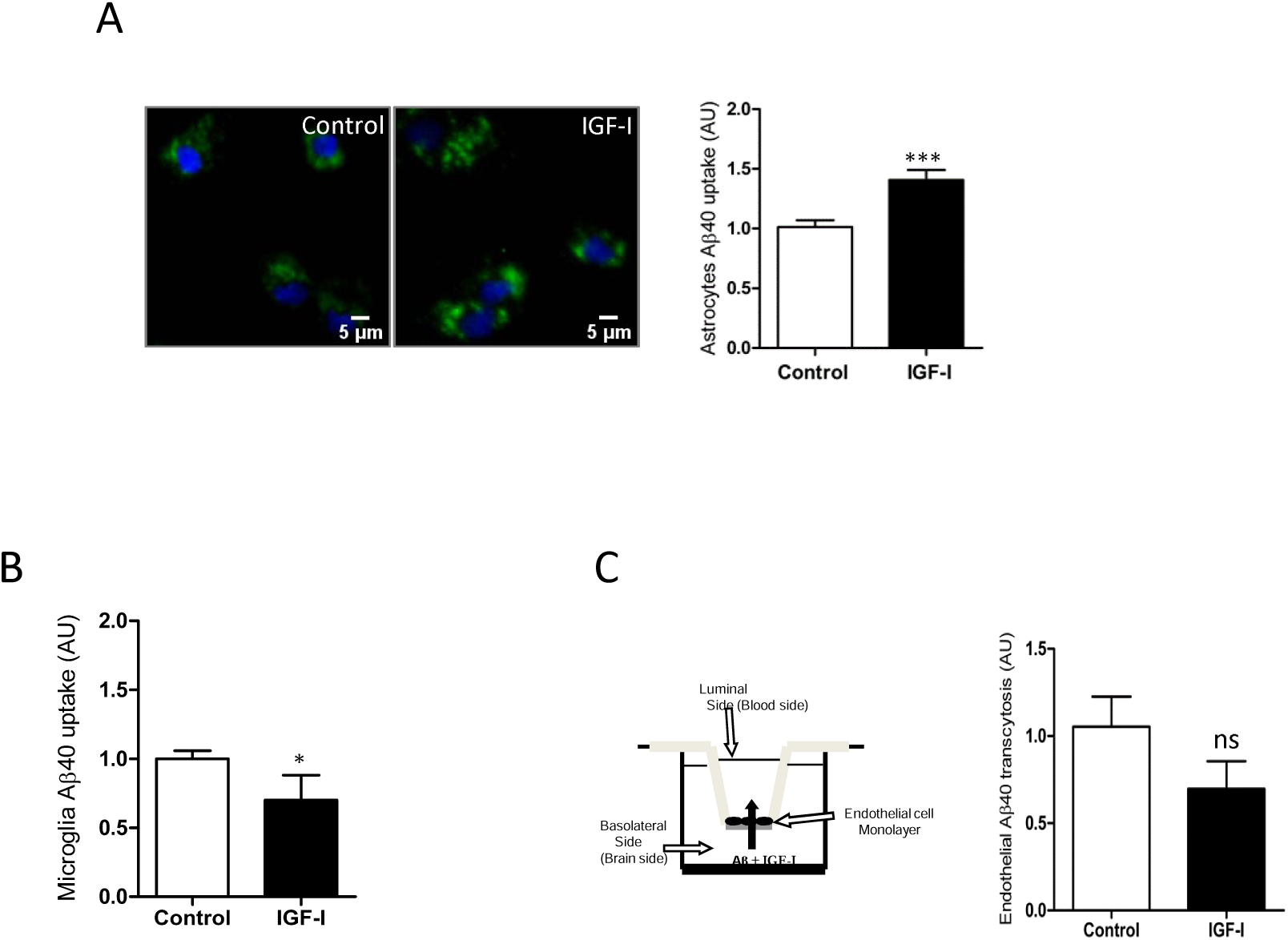
IGF-I modulates brain Aβ uptake in a cell-specific manner. **A**, Aβ uptake by astrocytes is significantly increased by IGF-I (n= 8). Representative photomicrograph showing uptake by cultured astrocytes of fluorescently labeled Aβ (green). Cell nuclei stained with Hoescht. **B**, Aβ uptake by microglia is significantly reduced by IGF-I (n= 7). **C**, IGF-I did not significantly affect brain-to-blood efflux in an in vitro system mimicking the blood-brain-barrier (cartoon in the left). Amount of Aβ in the upper chamber was quantified 15 h after adding it to the lower chamber in the presence or absence of IGF-I (n= 6).

### Reduced brain IGF-I activity in overweight mice

Since serum IGF-I levels are increased in overweight mice (Figure 2A), we determined whether brain IGF-I is correspondingly higher, as serum IGF-I crosses the BBB (16). However, overweight mice showed normal brain IGF-I levels (Figure 5A). To explain this discrepancy between peripheral and central IGF-I levels, we determined whether passage of serum IGF-I into the brain is reduced in overweight mice. To this end, we took advantage that exposure to environmental enrichment (EE) stimulates the passage of serum IGF-I into the brain (17). We tested whether overweight mice show altered passage of IGF-I after EE by measuring Tyr-phosphorylation of brain IGF-I receptors as a proxy of their activity. After EE, overweight mice showed reduced IGF-IR phosphorylation (Figure 5B), pointing to impaired entrance of circulating IGF-I. In addition, since systemic administration of IGF-I to lean mice housed under standard conditions showed enhanced APP/IGF-IR interaction (Figure 5C), corroborating in vitro observations (Figure 3D), we used this interaction as an additional indicator of the entrance of IGF-I into the brain of EE-stimulated overweight mice. Significantly, whereas in lean mice EE produced enhanced brain APP/IGF-IR interactions, in overweight mice, this interaction was significantly smaller (Figure 5D), pointing to reduced entrance of IGF-I.

**Figure 5.**
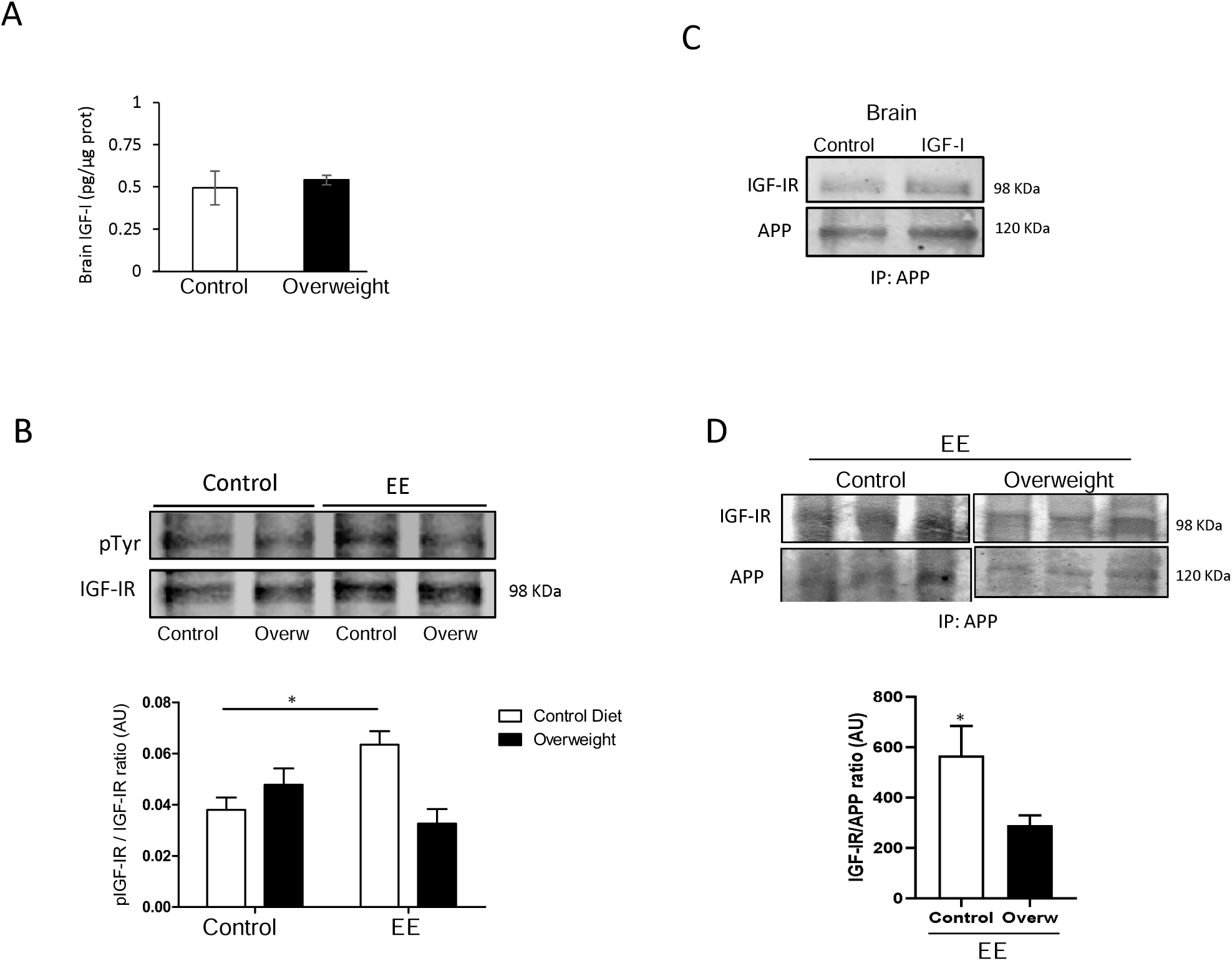
Reduced entrance of serum IGF-I in overweight mice. **A**, Brain levels of IGF-I were normal in HFD-fed overweight mice (n= 10 per group). **B**, In response to environmental enrichment (EE), overweight mice show lower brain IGF-IR phosphorylation than lean mice receiving a standard diet. Representative blot is shown, together with quantification histograms (n= 10 EE/ 6 Control; for each diet). **C**, Brain IGF-IR/APP co-immunoprecipitation is increased after systemic IGF-I administration. **D**, Interaction of APP with IGF-IR in the brain of mice submitted to EE stimulation was significantly decreased in overweight mice (n=10 per group).

## Discussion

These results indicate that diet influences peripheral Aβ clearance through the liver without impacting brain Aβ levels. A lack of correlation between peripheral and central Aβ clearance agrees with observations that reducing peripheral Aβ does not affect brain Aβ levels, or with the absence of correlation between central and peripheral Aβ levels in AD patients (36-39). However, increased peripheral Aβ levels after anti-Aβ treatment was reported to parallel a decrease of brain Aβ; reducing peripheral Aβ was sufficient to reduced brain Aβ, and recent studies favor a diagnostic utility of the relationship between plasma and CSF Aβ_1-42_ (27,40,41). Thus, the relationship between peripheral and central Aβ is still under debate (42); indeed, a substantial part of brain Aβ clearance in humans takes place in the periphery (43). In turn, normal brain levels of Aβ in overweight mice agree with previously reported similar observations (44), but not with increased brain Aβ load found by others (45,46). Conversely, enhanced elimination of circulating Aβ in overweight mice favors the notion that higher body mass index may be protective rather than detrimental for AD risk (4), However, unaffected brain Aβ load does not fit with a protective effect of increased body mass, unless still undefined systemic changes contribute to AD, as recently postulated (47).

Among them, we considered circulating IGF-I as a probable systemic factor influencing the connection of obesity with AD. IGF-I is involved in brain Aβ clearance (12) -although this has been questioned (48), and shows diet-sensitive actions in the brain (14). Indeed, several observations favor the involvement of circulating IGF-I in the systemic pro-clearance actions of a high fat diet. 1) IGF-I levels are increased in overweight mice, 2) IGF-I, as previously seen with insulin (26), stimulates uptake of Aβ by hepatocytes, and 3) LID mice with low serum IGF-I show reduced peripheral Aβ clearance. Thus, higher serum IGF-I levels in overweight mice may contribute to reduce peripheral Aβ levels.

As indicated by an increased sAPPα/sAPPβ ratio in IGF-I-treated neurons and astrocytes, the net action of IGF-I on the main cell types producing Aβ in the brain is to favor non-amyloidogenic processing of APP, contributing in this way to lower its brain levels and enhance neuroprotection, as sAPPα is neuroprotective acting in part through IGF-IR (49). Thus, the overall action of IGF-I in the brain may be anti-amyloidogenic. Intriguingly, insulin favors Aβ secretion in neurons (50), suggesting a complex interplay of these hormones in regulating brain Aβ levels. At the same time, reduced IGF-I entrance in the brain of overweight mice may hamper its anti-amyloidogenic actions. Indeed, overweight mice showed not only reduced entrance of serum IGF-I in response to EE stimulation, as determined by reduced brain IGF-IR phosphorylation, but also reduced APP/IGF-IR interaction. In previous work we documented an inhibitory effect of triglycerides (TGLs) in BBB entrance of IGF-I across the choroid plexus (14). It is possible that high serum TGLs as a result of the high fat diet interfere also with the BBB entrance of IGF-I across brain endothelial cells in overweight mice.

Reduced IGF-I entrance would affect its pro-clearance actions on brain Aβ (12,51). Also, we cannot discard that the inhibitory actions of IGF-I on Aβ uptake by microglia may also counteract its actions on astrocytes (but see below). Alternatively, brain Aβ levels may not be affected by peripheral Aβ clearance or other factors may also contribute to it, such as the recently postulated vascular drainage (52,53). Interestingly, insulin also enhances the degradation of Aβ and its clearance in astrocytes (54), and hepatocytes (26), respectively. Thus, these two closely related hormones may modulate Aβ disposal in a concerted manner, as previously reported for glucose handling (55).

IGF-I stimulates Aβ uptake by astrocytes, while inhibits it in microglia. Whereas astrocytes appear critical to determine Aβ load (56), and increased clearance of Aβ by astrocytes may result in reduced Aβ plaques (34), inhibition of Aβ uptake by microglia may also reduce plaques (57), as the role of microglial uptake of Aβ in plaque formation may be detrimental (58,59). In accordance with a stimulatory effect of IGF-I on astrocytes, previous observations suggested that astrocyte-derived IGF-I protects neurons against Aβ toxicity through a mechanism involving its uptake (60).

The observed astrocyte-specific interaction of APP with IGF-IR and on sAPPα and sAPPβ levels may be related to a differential processing of APP by IGF-I in these cells, since its processing depends on its intracellular localization (31). Of the different isoforms of APP, the major one expressed in neurons is APP_695_, that lacks the extracelular protein-protein interaction domain KPI. This domain is present in the longer isoforms, APP_751_ and APP_770_, that are the most abundant types in glial cells (61). It is possible that KPI is involved in the observed interactions with IGF-IR in astrocytes. In turn, a trend of IGF-I to inhibit brain efflux of Aβ through BBB endothelial cells, would favor its accumulation in brain parenchyma (62). We previously reported that IGF-I stimulates Aβ efflux through the choroid plexus BBB (12), an observation supported by the reducing effects of in vivo IGF-I administration on brain Aβ levels (12,51). Thus, IGF-I may show site-specific effects on Aβ efflux through BBB cells.

An important limitation of this study is that we determined peripheral Aβ clearance indirectly. Measuring circulating levels of Aβ in overweight mice would be necessary to firmly establish that peripheral Aβ clearance is enhanced. However, available methods of quantification of serum Aβ are not sensitive enough to reliably detect decreases in non-transgenic mice.

In summary, diet influences peripheral, but not central Aβ clearance. A lack of correlation between peripheral clearance and central Aβ in overweight mice further support a non-linear relationship between both compartments. Actions of IGF-I on Aβ handling may be relevant in AD pathology and related to diet influences on the disease; therefore cellular sites of IGF-I interaction may constitute new druggable targets, through, for example, potentiation of IGF-I-induced hepatic Aβ clearance.

## Materials and Methods

### Materials

Human IGF-I was purchased from PeProTec (UK). Primary antibodies were monoclonal anti-IGF-I receptor (1:1000; Santa Cruz Biotechnology, USA), monoclonal anti-APP (Nt 22C11; Millipore; 1:200), for PLA studies, polyclonal anti-APP (Sigma; 1:200), for immunoprecipitation, and monoclonal anti-pTyr (1:1000, Transduction Labs, USA). Secondary antibodies were goat anti-rabbit (1:20000) or mouse IRDye-coupled (1:20000), both from LI-COR (USA).

### Animals

Male adult (3-5 months old) and new-born wild type C57BL6/J mice, and adult liver IGF-I deficient mutant mice (LID mice; bred in-house, congenic with C57/BL6/J) were used. LID mice present low levels of serum IGF-I due to the disruption of the liver IGF-I gene with the albumin-Cre/Lox system (24). Serum IGF-I deficient mice have normal body and brain weights and they do not show any major developmental defects (24,63). Animal procedures followed European (86/609/EEC & 2003/65/EC, European Council Directives) and approval of the local Bioethics Committee.

### High fat diet

Wild type C57BL6/J mice were fed for 10 weeks with either a control diet (ref E15000-04), or a high fat diet (HFD) with 45% KJ fat + 1.25% cholesterol (ref E15744-34), both purchased from ssniff Spezialdiäten GmbH (Germany). After 10 weeks, animals were overweighed (Suppl Fig A), developed glucose intolerance (Supp Fig B), together with hyperinsulinemia and insulin resistance (not shown).

### Cell cultures

Astroglial cultures with >95% GFAP-positive cells were prepared as described (64). Postnatal (day 1-2) brains were dissected, forebrain removed, and mechanically dissociated. The resulting mixed cell suspension was centrifuged and plated in DMEM/F-12 (Life Technologies) with 10% fetal bovine serum (Life Technologies) and 100 mg/ml of antibiotic-antimycotic solution (Sigma-Aldrich, Spain). When confluent, cells were shaken (210 rpm/37°C/3 hours) to detach microglial cells. For microglial cultures, supernatants were centrifuged (1000 rpm/5 min), re-suspended in DMEM/F12 (Life Technologies)+FBS (Gibco, USA), HS and penicillin/streptomycin solution. Cells were seeded at 12.5-10^4^ cells/cm^2^ in a multi-well coated with poly-L-lysine (65), and cultured for 2 days. Cells were then changed to DMEM/F12 for 3 hours until Aβ uptake was carried out (see below). Astrocytes were then collected from the same flasks that microglia was obtained, as follows. After removing the microglia-containing supernatant, the medium was replaced and flasks shaken for 15 h/280 rpm. Cells were then trypsinized and seeded at 3.75×10^4^ cells /cm^2^ in the same culture medium, replaced every 4 days. When 80% confluency was reached, astrocytes were cultured for 3 hours with DMEM/F12 before the different assays were initiated (see below). Endothelial cell cultures were performed as described (65). Briefly, dissection was performed on ice and cortices were cut into small pieces (1 mm^3^), digested in a mixture of collagenase/dispase (270 U collagenase/ml, 10% dispase) and DNAse (10 U/ml) in DMEM for 1.5 h at 37°C. The cell pellet was separated by centrifugation in 20% bovine serum albumin/DMEM (1000g, 5 min). Capillary fragments were retained on a 10µm nylon filter, removed from the filter with endothelial cell basal medium (Life Technologies), supplemented with 20% bovine plasma-derived serum and antibiotics (penicillin, 100 U/ml; streptomycin, 100 µg/ml), and seeded on 60-mm Petri dishes multi-well plate coated with collagen type IV (5 µg/cm^2^) and fibronectin (1µg/cm^2^). 3µg/mL puromycin was added for 3 days, removed from the culture medium and replaced by fibroblast growth factor (2 ng/ml) and hydrocortisone (1µg /ml). For hepatocytes cultures, adult (2 months old) control animals were anesthesized (pentobarbital 50 mg/kg), and the hepatic portal vein exposed to inject a solution containing NaCl (118 Mm), KCl (4.7 Mm), KH_2_PO_4_ (1.2 Mm), NaHCO_3_ (25Mm), glucose (5.5 Mm), and EGTA (0.5 Mm) at 37C. The inferior cava vein was cut to open the circuit. Thereafter the same solution without EGTA and containing CaCl_2_ (2 Mm), MgSO_4_ (1.2 Mm), and colagenase (90U/ml) was perfused. The liver was dissected and placed in DMEM/F12 -10% FBS with penicillin/ streptomycin, filtered in a 70 um Nylon mesh, centrifuged (60g, 5 min) and re-suspended in DMEM/F12-10% FBS with 45% Percoll® (Sigma Aldrich). Cells were then re-suspended and washed 3X in DMEM/F12-10% FBS using 200g, 10 min spins, before plating them at 8.25 x 10^4^ cells/collagen-coated multi-well. Cultures were kept 2 days before use.

### Glucose tolerance test (GTT)

Mice were fasted for 6 hours and left isolated in individual cages (with water but no food access) for at least 30 min before starting the test to avoid any stress-related effect on glycemia (66). For the glucose tolerance test (GTT), an overload of glucose (2g/kg) was injected intraperitoneally. The aqueous solution was left overnight at room temperature so the β-form of glucose was enriched. Blood samples were extracted from the tip of the tail at time 0, 15, 30, 60, and 90 to measure glucose levels with a glucometer (Menarini Diagnosis, Italy).

### Environmental enrichment

Mice were submitted to environmental enrichment as explained in detail elsewhere (67). Briefly, animals were placed for 2 hours in a large cage, 10 animals/cage and with different objects (cardboard tunnels, shelters of different materials, a plastic net, toys, chewable and nesting material). Thereafter, they were sacrificed and their brain collected for immunoprecipitation and western blot analysis.

### Aβ uptake

*In vitro*: Cells were treated during 15 hours with 500 nM soluble Aβ40-HiLyte Fluor™ 488 (AnaSpec) (68), and IGF-I (1 nM in glial cultures, 10 nM in hepatocytes). Thereafter, cultures were washed with PBS pH 6.0 to eliminate membrane bound Aβ followed by PBS pH 7.4. Cell nuclei were stained with Hoechst 33342 (Thermo Fisher Scientific; 1:500) in PBS pH 7.4/5 min, fluorescent images were taken in an DMI 6000 (Leica) microscope using Exc: 350 nm/ Em: 461 nm for Hoeschst dye and Exc: 503 nm/ Em: 528 nm for fluorescently labeled Aβ. Thereafter, cells were lysed in Tris-HCl (10 mM) pH 8.0, guanidine (50 mM), and spinned at 14.000 rpm for 10 min at 4°C. Fluorescence was quantified in a FLUOStar OPTIMA (BMG Labtech) at Exc: 485 nm/ Em: 520 nm. In transcytosis assays using brain endothelial cells, Aβ40-HiLyte Fluor™ 488 soluble (500 nM) was added in the bottom compartment (Figure 4C) with or without 1 nM IGF-I, and after 15 hours the culture medium from the upper chamber was collected and fluorescence measured in the fluorimeter, as above. *In vivo*: Aβ40-HiLyte Fluor™ 488 (400 μg/kg) was injected into the tail vein using a 0.38 mm cannula (Intramedic, Spain), and after 90 min mice were sacrificed, blood taken from the heart and liver dissected. Liver tissue was homogenized in Tris-HCl (10 mM) pH 8.0 - guanidine (50 mM). Fluorescence in serum and liver extracts was quantified by fluorimetry, as above. Values were normalized per ml of serum or mg of protein. The latter was measured in liver samples using the BCA system (Sigma).

### Immunoassays

Western blot and immunoprecipitation were performed as described elsewhere in detail (69). Densitometric analysis of blots was performed using the Odissey system (Lycor Biosciences, USA). A representative blot is shown from a total of at least three independent experiments. GFAP immunocytochemistry in cultured cells followed previously published procedures (69). In brief, cultured cells were incubated to block non-specific antibody binding, followed by incubation overnight at 4°C with anti-GFAP in phosphate buffer (PB) - 1% bovine albumin - 1% Triton X-100 (PBT). After several washes in PB, sections were incubated with an Alexa-coupled secondary antibody (1:1000, Molecular Probes, USA) diluted in PBT. Finally, a 1:500 dilution (in PBS) of DAPI (Hoechst 33342) was added for 3 minutes. Wells were rinsed several times in PB 0.1 N, mounted with 15 µl of gerbatol mounting medium, and allowed to dry. Omission of primary antibody was used as control. Microphotographs were taken in a Leica (Germany) microscope. Plaque load was determined as explained elsewhere in detail (69).

IGF-I in serum and brain was determined using a species-specific ELISA (R&D Systems, USA), as described in detail elsewhere (18). Murine Aβ (Thermofisher, USA), and murine sAPPα and sAPPβ were determined by ELISA in brain lysates and culture supernatants, respectively, following the manufacturer’s instructions. Blood was collected from the heart after pentobarbital anesthesia and thereafter brains were dissected and frozen at -80°C until used.

### Proximity ligation assays (PLA)

Assays were performed as described (70). Amyloid precursor protein (APP) – IGF-IR interactions were detected in astrocytes and neurons grown on glass coverslips using the Duolink II in situ PLA detection Kit (OLink; Bioscience, Sweden). Cultured cells were fixed in 4% paraformaldehyde/10 min, washed with PBS containing 20 mM glycine to quench the aldehyde groups, permeabilized with the same buffer containing 0.05% Triton X-100 for 5 min, and washed with PBS. After 1 h/37°C with the blocking solution in a pre-heated humidity chamber, cells were incubated overnight in antibody diluent medium with primary antibodies: mouse monoclonal anti-APP and rabbit polyclonal anti-IGF-I receptor, and processed following the instructions of the supplier using the PLA probes detecting rabbit or mouse antibodies (Duolink II PLA probe anti-Rabbit plus, and Duolink II PLA probe anti-Mouse minus, diluted 1:5 in antibody diluent), and a DAPI-containing mounting medium.

### Statistical analysis

Normal distribution tests were carried out in all experiments and a non-parametric Wilcoxon test was applied accordingly. For samples with normal distribution, parametric tests include one-way ANOVA followed by a Bonferroni or t-test. A p<0.05 was considered significant.

## Acknowledgements

We are thankful to M. Garcia for technical support. This work was funded by a grant from Ciberned, by an Inter-CIBER project (PIE14/00061), and from SAF2013-40710-R (AEI/FEDER, UE).

## Authors contributions

RH performed experiments, analyzed results, and wrote part of the manuscript. ATS performed experiments and analyzed results. LMR performed experiments and analyzed results. EFS performed experiments and analyzed results. SDP helped with experiments. AMF performed experiments and analyzed results. ITA designed the study, analyzed results, and wrote the manuscript.

## LEGENDS TO FIGURES

**Supplementary Figure. A**, Mice submitted to a high fat diet during 10 weeks (W) show increased body weight along time (n=10 per group). **B**, Overweight mice show reduced tolerance to systemic glucose load, as determined by significantly greater blood glucose levels over 90 min after glucose administration.

## References

1. Kivimaki M, Luukkonen R, Batty GD, Ferrie JE, Pentti J, Nyberg ST, Shipley MJ, Alfredsson L, Fransson EI, Goldberg M, Knutsson A, Koskenvuo M, Kuosma E, Nordin M, Suominen SB, Theorell T, Vuoksimaa E, Westerholm P, Westerlund H, Zins M, Kivipelto M, Vahtera J, Kaprio J, Singh-Manoux A, Jokela M. Body mass index and risk of dementia: Analysis of individual-level data from 1.3 million individuals. Alzheimers Dement 2018; 14:601–609

2. Profenno LA, Porsteinsson AP, Faraone SV. Meta-analysis of Alzheimer’s disease risk with obesity, diabetes, and related disorders. Biol Psychiatry 2010; 67:505–512

3. Singh-Manoux A, Dugravot A, Shipley M, Brunner EJ, Elbaz A, Sabia S, Kivimaki M. Obesity trajectories and risk of dementia: 28 years of follow-up in the Whitehall II Study. Alzheimers Dement 2018; 14:178–186

4. Lee CM, Woodward M, Batty GD, Beiser AS, Bell S, Berr C, Bjertness E, Chalmers J, Clarke R, Dartigues J-F, Davis-Plourde K, Debette S, Di Angelantonio E, Feart C, Frikke-Schmidt R, Gregson J, Haan MN, Hassing LB, Hayden KM, Hoevenaar-Blom MP, Kaprio J, Kivimaki M, Lappas G, Larson EB, LeBlanc ES, Lee A, Lui L-Y, Moll van Charante EP, Ninomiya T, Nordestgaard LT, Ohara T, Ohkuma T, Palviainen T, Peres K, Peters R, Qizilbash N, Richard E, Rosengren A, Seshadri S, Shipley M, Singh-Manoux A, Strand BH, van Gool WA, Vuoksimaa E, Yaffe K, Huxley RR. Association of anthropometry and weight change with risk of dementia and its major subtypes: A meta-analysis consisting 2.8 million adults with 57 294 cases of dementia. Obes Rev 2020:10.1111/obr.12989

5. Dye L, Boyle NB, Champ C, Lawton C. The relationship between obesity and cognitive health and decline. Proc Nutr Soc 2017; 76:443–454

6. Jimenez A, Pegueroles J, Carmona-Iragui M, Vilaplana E, Montal V, Alcolea D, Videla L, Illan-Gala I, Pane A, Casajoana A, Belbin O, Clarimon J, Moize V, Vidal J, Lleo A, Fortea J, Blesa R, Alzheimer’s Disease Neuroimaging I. Weight loss in the healthy elderly might be a non-cognitive sign of preclinical Alzheimer’s disease. Oncotarget 2017; 8:104706–104716

7. Sun Z, Wang Z-T, Sun F-R, Shen X-N, Xu W, Ma Y-H, Dong Q, Tan L, Yu J-T, Alzheimer’s Disease Neuroimaging I. Late-life obesity is a protective factor for prodromal Alzheimer’s disease: a longitudinal study. Aging (Albany NY) 2020; 12:2005–2017

8. Scheltens P, Blennow K, Breteler MM, de Strooper B, Frisoni GB, Salloway S, Van der Flier WM. Alzheimer’s disease. Lancet 2016; 388:505–517

9. Engin A. The Definition and Prevalence of Obesity and Metabolic Syndrome. Adv Exp Med Biol 2017; 960:1–17

10. Cox AJ, West NP, Cripps AW. Obesity, inflammation, and the gut microbiota. Lancet Diabetes Endocrinol 2015; 3:207–215

11. Fernandez AM, Santi A, Torres Aleman I. Insulin Peptides as Mediators of the Impact of Life Style in Alzheimer’s disease. Brain plasticity (Amsterdam, Netherlands) 2018; 4:3–15

12. Carro E, Trejo JL, Gomez-Isla T, LeRoith D, Torres-Aleman I. Serum insulin-like growth factor I regulates brain amyloid-beta levels. Nat Med 2002; 8:1390–1397

13. Cohen E, Paulsson JF, Blinder P, Burstyn-Cohen T, Du D, Estepa G, Adame A, Pham HM, Holzenberger M, Kelly JW, Masliah E, Dillin A. Reduced IGF-1 signaling delays age-associated proteotoxicity in mice. Cell 2009; 139:1157–1169

14. Dietrich MO, Muller A, Bolos M, Carro E, Perry ML, Portela LV, Souza DO, Torres-Aleman I. Western Style Diet Impairs Entrance of Blood-Borne Insulin-like Growth Factor-1 into the Brain. Neuromolecular Med 2007; 9:324–330

15. Bach MA, Shen-Orr Z, Lowe WL, Jr., Roberts CT, Jr., LeRoith D. Insulin-like growth factor I mRNA levels are developmentally regulated in specific regions of the rat brain. Brain Res Mol Brain Res 1991; 10:43–48

16. Carro E, Nunez A, Busiguina S, Torres-Aleman I. Circulating insulin-like growth factor I mediates effects of exercise on the brain. J Neurosci 2000; 20:2926–2933

17. Nishijima T, Piriz J, Duflot S, Fernandez AM, Gaitan G, Gomez-Pinedo U, Verdugo JM, Leroy F, Soya H, Nunez A, Torres-Aleman I. Neuronal activity drives localized blood-brain-barrier transport of serum insulin-like growth factor-I into the CNS. Neuron 2010; 67:834–846

18. Trejo JL, Piriz J, Llorens-Martin MV, Fernandez AM, Bolos M, LeRoith D, Nunez A, Torres-Aleman I. Central actions of liver-derived insulin-like growth factor I underlying its pro-cognitive effects. Mol Psychiatry 2007; 12:1118–1128

19. Jacobsen KT, Adlerz L, Multhaup G, Iverfeldt K. Insulin-like growth factor-1 (IGF-1)-induced processing of amyloid-{beta} precursor protein (APP) and APP-like protein 2 is mediated by different metalloproteinases. J Biol Chem 2010; 285:10223–10231

20. Adlerz L, Holback S, Multhaup G, Iverfeldt K. IGF-1-induced processing of the amyloid precursor protein family is mediated by different signaling pathways. J Biol Chem 2007; 282:10203–10209

21. Zhang H, Gao Y, Dai Z, Meng T, Tu S, Yan Y. IGF-1 reduces BACE-1 expression in PC12 cells via activation of PI3-K/Akt and MAPK/ERK1/2 signaling pathways. Neurochem Res 2011; 36:49–57

22. Araki W, Kume H, Oda A, Tamaoka A, Kametani F. IGF-1 promotes beta-amyloid production by a secretase-independent mechanism. Biochem Biophys Res Commun 2009;

23. Costantini C, Scrable H, Puglielli L. An aging pathway controls the TrkA to p75NTR receptor switch and amyloid beta-peptide generation. EMBO J 2006; 25:1997–2006

24. Yakar S, Liu JL, Stannard B, Butler A, Accili D, Sauer B, LeRoith D. Normal growth and development in the absence of hepatic insulin-like growth factor I. Proc Natl Acad Sci U S A 1999; 96:7324–7329

25. Ghiso J, Shayo M, Calero M, Ng D, Tomidokoro Y, Gandy S, Rostagno A, Frangione B. Systemic Catabolism of Alzheimer’s A{beta}40 and A{beta}42. Journal of Biological Chemistry 2004; 279:45897–45908

26. Tamaki C, Ohtsuki S, Terasaki T. Insulin facilitates the hepatic clearance of plasma amyloid beta-peptide (1 40) by intracellular translocation of low-density lipoprotein receptor-related protein 1 (LRP-1) to the plasma membrane in hepatocytes. Mol Pharmacol 2007; 72:850–855

27. DeMattos RB, Bales KR, Cummins DJ, Dodart JC, Paul SM, Holtzman DM. Peripheral anti-A beta antibody alters CNS and plasma A beta clearance and decreases brain A beta burden in a mouse model of Alzheimer’s disease. Proc Natl Acad Sci U S A 2001; 98:8850–8855

28. Zegarra-Valdivia JA, Santi A, Fernandez de Sevilla ME, Nunez A, Torres Aleman I. Serum Insulin-Like Growth Factor I Deficiency Associates to Alzheimer’s Disease Co-Morbidities. J Alzheimers Dis 2019; 69:979–987

29. Kamenetz F, Tomita T, Hsieh H, Seabrook G, Borchelt D, Iwatsubo T, Sisodia S, Malinow R. APP processing and synaptic function. Neuron 2003; 37:925–937

30. Zhao J, O’Connor T, Vassar R. The contribution of activated astrocytes to Abeta production: implications for Alzheimer’s disease pathogenesis. J Neuroinflammation 2011; 8:150

31. Choy RW, Cheng Z, Schekman R. Amyloid precursor protein (APP) traffics from the cell surface via endosomes for amyloid beta (Abeta) production in the trans-Golgi network. Proc Natl Acad Sci U S A 2012; 109:E2077–E2082

32. Pietrzik CU, Yoon IS, Jaeger S, Busse T, Weggen S, Koo EH. FE65 constitutes the functional link between the low-density lipoprotein receptor-related protein and the amyloid precursor protein. J Neurosci 2004; 24:4259–4265

33. Griciuc A, Serrano-Pozo A, Parrado AR, Lesinski AN, Asselin CN, Mullin K, Hooli B, Choi SH, Hyman BT, Tanzi RE. Alzheimer’s Disease Risk Gene CD33 Inhibits Microglial Uptake of Amyloid Beta. Neuron 2013;

34. Wyss-Coray T, Loike JD, Brionne TC, Lu E, Anankov R, Yan F, Silverstein SC, Husemann J. Adult mouse astrocytes degrade amyloid-beta in vitro and in situ. Nat Med 2003; 9:453–457

35. Deane R, Wu Z, Sagare A, Davis J, Du YS, Hamm K, Xu F, Parisi M, LaRue B, Hu HW, Spijkers P, Guo H, Song X, Lenting PJ, Van Nostrand WE, Zlokovic BV. LRP/amyloid beta-peptide interaction mediates differential brain efflux of Abeta isoforms. Neuron 2004; 43:333–344

36. Walker JR, Pacoma R, Watson J, Ou W, Alves J, Mason DE, Peters EC, Urbina HD, Welzel G, Althage A, Liu B, Tuntland T, Jacobson LH, Harris JL, Schumacher AM. Enhanced Proteolytic Clearance of Plasma A+¦ by Peripherally Administered Neprilysin Does Not Result in Reduced Levels of Brain A+¦ in Mice. The Journal of Neuroscience 2013; 33:2457–2464

37. Fukumoto H, Tennis M, Locascio JJ, Hyman BT, Growdon JH, Irizarry MC. Age but not diagnosis is the main predictor of plasma amyloid beta-protein levels. Arch Neurol 2003; 60:958–964

38. Mehta PD, Pirttila T, Patrick BA, Barshatzky M, Mehta SP. Amyloid beta protein 1-40 and 1-42 levels in matched cerebrospinal fluid and plasma from patients with Alzheimer disease. Neurosci Lett 2001; 304:102–106

39. Siemers ER, Dean RA, Friedrich S, Ferguson-Sells L, Gonzales C, Farlow MR, May PC. Safety, tolerability, and effects on plasma and cerebrospinal fluid amyloid-beta after inhibition of gamma-secretase. Clin Neuropharmacol 2007; 30:317–325

40. Albani D, Marizzoni M, Ferrari C, Fusco F, Boeri L, Raimondi I, Jovicich J, Babiloni C, Soricelli A, Lizio R, Galluzzi S, Cavaliere L, Didic M, Schonknecht P, Molinuevo JL, Nobili F, Parnetti L, Payoux P, Bocchio L, Salvatore M, Rossini PM, Tsolaki M, Visser PJ, Richardson JC, Wiltfang J, Bordet R, Blin O, Forloni G, Frisoni GB, PharmaCog C. Plasma Abeta42 as a Biomarker of Prodromal Alzheimer’s Disease Progression in Patients with Amnestic Mild Cognitive Impairment: Evidence from the PharmaCog/E-ADNI Study. J Alzheimers Dis 2019; 69:37–48

41. Sutcliffe JG, Hedlund PB, Thomas EA, Bloom FE, Hilbush BS. Peripheral reduction of beta-amyloid is sufficient to reduce brain beta-amyloid: implications for Alzheimer’s disease. J Neurosci Res 2011; 89:808–814

42. Bassendine MF, Taylor-Robinson SD, Fertleman M, Khan M, Neely D. Is Alzheimer’s Disease a Liver Disease of the Brain? Journal of Alzheimer’s Disease 2020; 75:1–14

43. Roberts KF, Elbert DL, Kasten TP, Patterson BW, Sigurdson WC, Connors RE, Ovod V, Munsell LY, Mawuenyega KG, Miller-Thomas MM, Moran CJ, Cross DT, 3rd, Derdeyn CP, Bateman RJ. Amyloid-beta efflux from the central nervous system into the plasma. Ann Neurol 2014; 76:837–844

44. Zhang L, Dasuri K, Fernandez-Kim SO, Bruce-Keller AJ, Freeman LR, Pepping JK, Beckett TL, Murphy MP, Keller JN. Prolonged diet induced obesity has minimal effects towards brain pathology in mouse model of cerebral amyloid angiopathy: implications for studying obesity-brain interactions in mice. Biochim Biophys Acta 2013; 1832:1456–1462

45. Li J, Deng J, Sheng W, Zuo Z. Metformin attenuates Alzheimer’s disease-like neuropathology in obese, leptin-resistant mice. Pharmacol Biochem Behav 2012; 101:564–574

46. Puig KL, Floden AM, Adhikari R, Golovko MY, Combs CK. Amyloid precursor protein and proinflammatory changes are regulated in brain and adipose tissue in a murine model of high fat diet-induced obesity. PLoS One 2012; 7:e30378

47. Wang J, Gu BJ, Masters CL, Wang Y-J. A systemic view of Alzheimer disease - insights from amyloid-β metabolism beyond the brain. Nature reviews Neurology 2017; 13:612–623

48. Lanz TA, Salatto CT, Semproni AR, Marconi M, Brown TM, Richter KE, Schmidt K, Nelson FR, Schachter JB. Peripheral elevation of IGF-1 fails to alter Abeta clearance in multiple in vivo models. Biochem Pharmacol 2008; 75:1093–1103

49. Jimenez S, Torres M, Vizuete M, Sanchez-Varo R, Sanchez-Mejias E, Trujillo-Estrada L, Carmona-Cuenca I, Caballero C, Ruano D, Gutierrez A, Vitorica J. Age-dependent accumulation of soluble Abeta oligomers reverses the neuroprotective effect of sAPPalpha by modulating PI3K/Akt-GSK-3beta pathway in Alzheimer mice model. J Biol Chem 2011;

50. Gasparini L, Gouras GK, Wang R, Gross RS, Beal MF, Greengard P, Xu H. Stimulation of beta-amyloid precursor protein trafficking by insulin reduces intraneuronal beta-amyloid and requires mitogen-activated protein kinase signaling. J Neurosci 2001; 21:2561–2570

51. Carro E, Trejo JL, Gerber A, Loetscher H, Torrado J, Metzger F, Torres-Aleman I. Therapeutic actions of insulin-like growth factor I on APP/PS2 mice with severe brain amyloidosis. Neurobiol Aging 2006; 27:1250–1257

52. van Veluw SJ, Hou SS, Calvo-Rodriguez M, Arbel-Ornath M, Snyder AC, Frosch MP, Greenberg SM, Bacskai BJ. Vasomotion as a Driving Force for Paravascular Clearance in the Awake Mouse Brain. Neuron 2019:S0896-6273(0819)30928-30926

53. Iliff JJ, Wang M, Liao Y, Plogg BA, Peng W, Gundersen GA, Benveniste H, Vates GE, Deane R, Goldman SA, Nagelhus EA, Nedergaard M. A Paravascular Pathway Facilitates CSF Flow Through the Brain Parenchyma and the Clearance of Interstitial Solutes, Including Amyloid +¦. Science Translational Medicine 2012; 4:147ra111–147ra111

54. Yamamoto N, Ishikuro R, Tanida M, Suzuki K, Ikeda-Matsuo Y, Sobue K. Insulin-signaling Pathway Regulates the Degradation of Amyloid beta-protein via Astrocytes. Neuroscience 2018; 385:227–236

55. Fernandez AM, Hernandez E, Guerrero-Gomez D, Miranda-Vizuete A, Torres Aleman I. A network of insulin peptides regulate glucose uptake by astrocytes: Potential new druggable targets for brain hypometabolism. Neuropharmacology 2018; 136:216–222

56. Katsouri L, Birch AM, Renziehausen AWJ, Zach C, Aman Y, Steeds H, Bonsu A, Palmer EOC, Mirzaei N, Ries M, Sastre M. Ablation of reactive astrocytes exacerbates disease pathology in a model of Alzheimer’s disease. Glia 2019:10.1002/glia.23759

57. Baik SH, Kang S, Son SM, Mook-Jung I. Microglia contributes to plaque growth by cell death due to uptake of amyloid beta in the brain of Alzheimer’s disease mouse model. Glia 2016; 64:2274–2290

58. Grathwohl SA, Kalin RE, Bolmont T, Prokop S, Winkelmann G, Kaeser SA, Odenthal J, Radde R, Eldh T, Gandy S, Aguzzi A, Staufenbiel M, Mathews PM, Wolburg H, Heppner FL, Jucker M. Formation and maintenance of Alzheimer’s disease [beta]-amyloid plaques in the absence of microglia. Nat Neurosci 2009; 12:1361–1363

59. Sosna J, Philipp S, Albay R, 3rd, Reyes-Ruiz JM, Baglietto-Vargas D, LaFerla FM, Glabe CG. Early long-term administration of the CSF1R inhibitor PLX3397 ablates microglia and reduces accumulation of intraneuronal amyloid, neuritic plaque deposition and pre-fibrillar oligomers in 5XFAD mouse model of Alzheimer’s disease. Mol Neurodegener 2018; 13:11

60. Pitt J, Wilcox KC, Tortelli V, Diniz LP, Oliveira MS, Dobbins C, Yu XW, Nandamuri S, Gomes FCA, DiNunno N, Viola KL, De Felice FG, Ferreira ST, Klein WL. Neuroprotective astrocyte-derived insulin/insulin-like growth factor 1 stimulates endocytic processing and extracellular release of neuron-bound Abeta oligomers. Mol Biol Cell 2017; 28:2623–2636

61. Belyaev ND, Kellett KA, Beckett C, Makova NZ, Revett TJ, Nalivaeva NN, Hooper NM, Turner AJ. The transcriptionally active amyloid precursor protein (APP) intracellular domain is preferentially produced from the 695 isoform of APP in a {beta}-secretase-dependent pathway. J Biol Chem 2010; 285:41443–41454

62. Jaeger LB, Dohgu S, Hwang MC, Farr SA, Murphy MP, Fleegal-DeMotta MA, Lynch JL, Robinson SM, Niehoff ML, Johnson SN, Kumar VB, Banks WA. Testing the neurovascular hypothesis of Alzheimer’s disease: LRP-1 antisense reduces blood-brain barrier clearance, increases brain levels of amyloid-beta protein, and impairs cognition. J Alzheimers Dis 2009; 17:553–570

63. Sjogren K, Jansson JO, Isaksson OG, Ohlsson C. A transgenic model to determine the physiological role of liver-derived insulin-like growth factor I. Minerva Endocrinol 2002; 27:299–311

64. Fernandez AM, Fernandez S, Carrero P, Garcia-Garcia M, Torres-Aleman I. Calcineurin in reactive astrocytes plays a key role in the interplay between proinflammatory and anti-inflammatory signals. J Neurosci 2007; 27:8745–8756

65. Trueba-Saiz A, Fernandez AM, Nishijima T, Mecha M, Santi A, Munive V, Torres-Aleman I. Circulating Insulin-like Growth Factor I Regulates Its Receptor in the Brain of Male Mice. Endocrinology 2017; 158:349–357

66. Ayala JE, Samuel VT, Morton GJ, Obici S, Croniger CM, Shulman GI, Wasserman DH, McGuinness OP, Consortium NIHMMPC. Standard operating procedures for describing and performing metabolic tests of glucose homeostasis in mice. Dis Model Mech 2010; 3:525–534

67. Trueba-Saiz A, Cavada C, Fernandez AM, Leon T, Gonzalez DA, Fortea OJ, Lleo A, Del ST, Nunez A, Torres-Aleman I. Loss of serum IGF-I input to the brain as an early biomarker of disease onset in Alzheimer mice. Transl Psychiatry 2013; 3:e330

68. Liu CC, Hu J, Zhao N, Wang J, Wang N, Cirrito JR, Kanekiyo T, Holtzman DM, Bu G. Astrocytic LRP1 Mediates Brain Abeta Clearance and Impacts Amyloid Deposition. J Neurosci 2017; 37:4023–4031

69. Fernandez AM, Jimenez S, Mecha M, Davila D, Guaza C, Vitorica J, Torres-Aleman I. Regulation of the phosphatase calcineurin by insulin-like growth factor I unveils a key role of astrocytes in Alzheimer’s pathology. Mol Psychiatry 2012; 17:705–718

70. Martinez-Rachadell L, Aguilera A, Perez-Domper P, Pignatelli J, Fernandez AM, Torres-Aleman I. Cell-specific expression of insulin/insulin-like growth factor-I receptor hybrids in the mouse brain. Growth Horm IGF Res 2019; 45:25–30

