## Supplementary figures and images for "DIET INFLUENCES PERIPHERAL AMYLOID β METABOLISM: A ROLE FOR CIRCULATING INSULIN-LIKE GROWTH FACTOR I"

### Suppl Figure

#### Slide 1
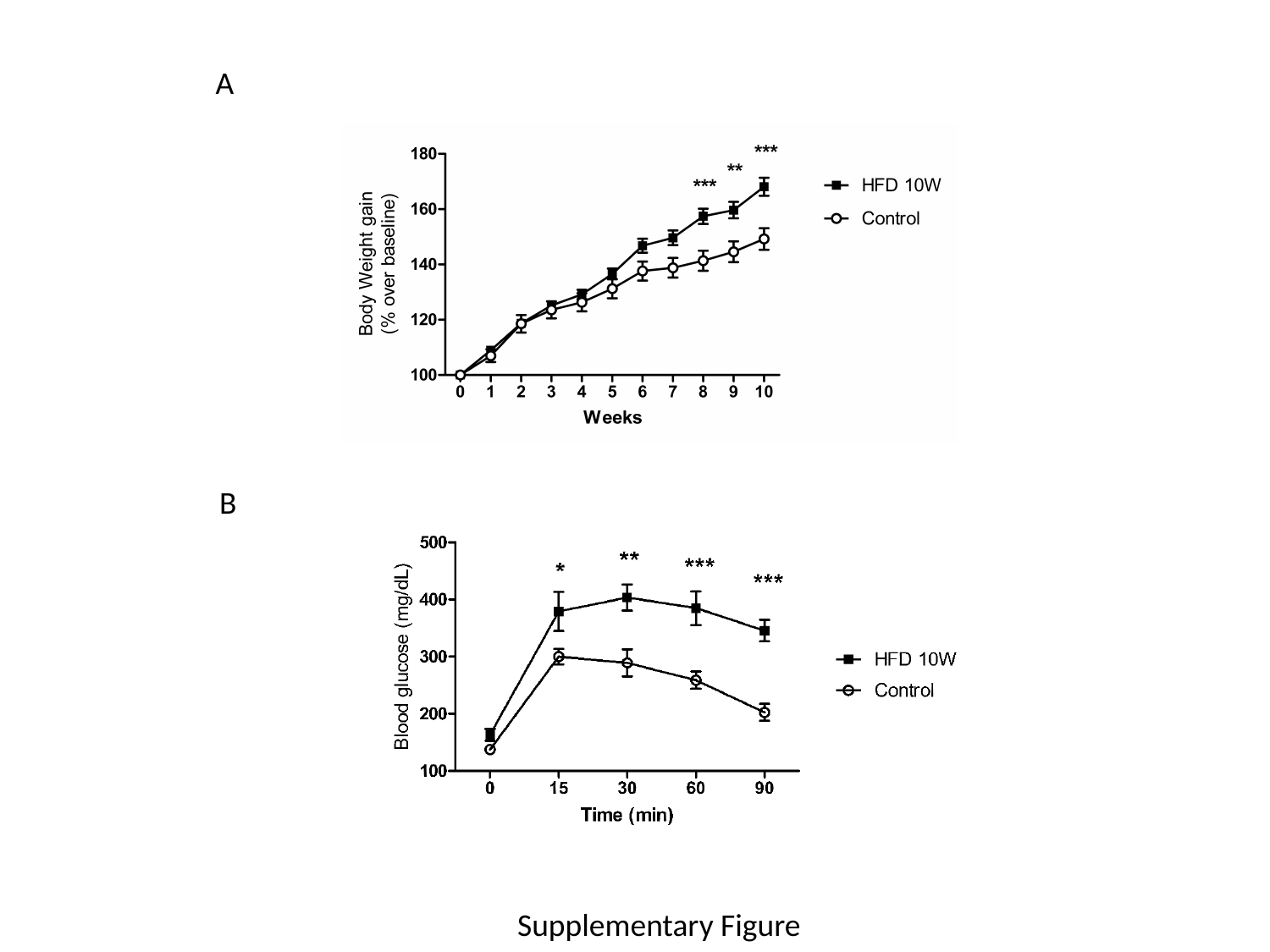

A
B
Supplementary Figure
